## supplementary codes for "A Stimulus-Computable Model for Audiovisual Perception and Spatial Orienting in Mammals": Alais_Burr_2004.m

```
%% Simulation of Alais & Burr (2004) using MCD and MLE models
% This script simulates the ventriloquist effect and compares predicted
% biases and JNDs from the MCD model and MLE predictions, using auditory
% and visual reliability manipulations. Human data is drawn from Figure 2
% of Alais & Burr (2004). Reproduces Figure 4C-E from Parise (2025) eLife.
% Alais & Burr (2004). The ventriloquist effect results from near-optimal bimodal integration. Current biology, 14(3), 257-262.
% 
% Author: Cesare Parise (2025)

clc; clear; close all;

%% Parameters
nPos = 500;       % Number of spatial samples
nOff = 5;         % Number of audiovisual disparities (offsets)
nSig = 3;         % Number of visual reliability levels
x = linspace(-50, 50, nPos);     % Spatial workspace
offset = linspace(-5, 5, nOff);  % AV disparity levels

sigA = 8.60;                   % Auditory JND (deg)
sigV = [1.25, 7.57, 19.78];    % Visual JNDs (deg)
sigA2 = sigA * ones(1, nSig);  % Duplicate for vector math

%% Initialize output matrices
biasMLE = nan(nSig, nOff);
sigmMLE = nan(nSig, nOff);
biasMCD = nan(nSig, nOff);
sigmMCD = nan(nSig, nOff);

%% Extracted human data from Alais & Burr (2004)
PSE_data = {
    [-4.99, -5.38; -2.50, -2.167; 0.02, 0.01; 2.48, 2.09; 4.92, 4.21],
    [-4.94, 0.01; -2.5, -0.60; -0.02, 0.49; 2.46, -0.29; 4.94, 0.36],
    [-4.94, 3.04; -2.48, 0.92; 0.02, 0.31; 2.46, -1.35; 4.92, -3.50]
};

%% Simulation
for ns = 1:nSig
    for no = 1:nOff
        % Define shifted unisensory stimuli
        A2 = normpdf(x, -offset(no), sigA); A2 = A2 / sum(A2);
        V2 = normpdf(x, +offset(no), sigV(ns)); V2 = V2 / sum(V2);
        
        %% MLE prediction
        wa = (1 / sigA^2) / (1 / sigA^2 + 1 / sigV(ns)^2);
        mumle = wa * (-offset(no)) + (1 - wa) * (+offset(no));
        sigmle = sqrt((sigA^2 * sigV(ns)^2) / (sigA^2 + sigV(ns)^2));
        
        biasMLE(ns, no) = mumle;
        sigmMLE(ns, no) = sigmle;

        %% MCD prediction 
        nSamp = 500;
        As = zeros(1, nSamp); As(10:150) = 1;
        Vs = zeros(1, nSamp); Vs(10:150) = 1;
        
        mcdAVxt = zeros(nSamp, nPos);
        for tt = 1:nPos
            v = Vs * V2(tt);
            a = As * A2(tt);
            signals = [v; a];
            [~, ~, MCD_output] = MCDq(signals, 500);
            mcdAVxt(:, tt) = sqrt(MCD_output.corr)';
        end

        mcd = sum(mcdAVxt); mcd = mcd / sum(mcd);

        % Fit Gaussian to MCD output (to get mu and sigma)
        param0 = [mumle, sigmle];
        fun = @(p) 1 - corr(normpdf(x, p(1), p(2))', mcd');
        options = optimset('MaxIter', 200, 'MaxFunEvals', 200);
        paramFit = fminsearch(fun, param0, options);

        biasMCD(ns, no) = paramFit(1);
        sigmMCD(ns, no) = paramFit(2);
    end
end

%% Plot: MCD vs human PSE
figure;
subplot(4,2,7); hold on; axis square;
colup = linspace(0,1,3);
for tt = 1:3
    col = [0,colup(tt), 1 - colup(tt)];
    plot(offset, biasMCD(tt,:), 'Color', col);
    scatter(PSE_data{tt}(:,1), PSE_data{tt}(:,2), 15, col, 'filled');
end
xlabel('Disparity (deg)'); ylabel('Bias (deg)');
title('MCD Prediction vs Human PSE');
xlim([-6 6]); ylim([-6 6]);

%% Plot: JND prediction vs human data
LocErrorAV = [4, 1.87; 32, 5.86; 64, 5.53];
LocErrorV  = [4, 1.25; 32, 7.57; 64, 19.78];
LocErrorA = 8.60;

subplot(4,2,8); hold on; axis square;
plot([0 65], [LocErrorA LocErrorA], 'r');
plot(LocErrorAV(:,1), LocErrorV(:,2), 'ob');
plot(LocErrorAV(:,1), sigmMLE(:,1), 'k');
plot(LocErrorAV(:,1), sigmMCD(:,1), 'ok');
scatter(LocErrorAV(:,1), LocErrorAV(:,2), 15, 'k', 'filled');
set(gca, 'YScale', 'log');
xlabel('Visual JND (deg)'); ylabel('Bimodal JND (deg)');
title('JND: Data vs Model Predictions');

%% Plot: Psychometric curves (MCD vs MLE)
xp = linspace(-20, 20, 100);
colup = linspace(0,1,5);

for tt = 1:3
    subplot(4,1,tt); hold on;
    for uu = 1:5
        yhatMCD = 1 - normcdf(xp, biasMCD(tt,uu), sigmMCD(tt,uu));
        yhatMLE = 1 - normcdf(xp, biasMLE(tt,uu), sigmMLE(tt,uu));
        col = [colup(uu), 1 - colup(uu), 0];
        plot(xp, yhatMCD, 'Color', col);
        plot(xp, yhatMLE, '.', 'Color', col, 'MarkerSize', 10);
    end
    xlim([-20 20]);
    xlabel('Probe Displacement (deg)'); ylabel('P(Left)');
    title(['MCD vs MLE: σ_V = ' num2str(sigV(tt)) ' deg']);
end
```
