## supplementary codes for "A Stimulus-Computable Model for Audiovisual Perception and Spatial Orienting in Mammals": MCD_Simulation_Alais_Carlile_2005.m

```
%% Simulation of Alais & Carlile (2005) using MCD model
% This script simulates the TOJ task for reverberabnt sounds
% Acoustic stimuli were grabbed from Alais & Carlile (2005) Figure 1
% TOJ data grabbed from Alais & Carlile (2005) Figure 2
% Reproduces Figure 2B from Parise (2025) eLife.
% Alais, D., & Carlile, S. (2005). Synchronizing to real events: Subjective audiovisual alignment scales with perceived auditory depth and speed of sound. Proceedings of the National Academy of Sciences, 102(6), 2244-2247.
% 
% Author: Cesare Parise (2025)
close all; clear; clc;

%% Load data
load data_Alais_Carlile
fs = data(1).fs;
duration1 = numel(data(1).signal) / fs;  % duration of signal in seconds
duration2 = 3;                           % total trial duration in seconds

% MCD model parameters: [alpha_V, alpha_A, tau_V, tau_A]
param = [0.0448821, 0.0369702, 0.180101, 0.180101];

%% Generate visual stimulus
vis = zeros(duration1 * fs, 1);
vis(round(0.019 * fs):round(0.032 * fs)) = 1;  % brief flash

st0 = zeros(duration2 * fs, 1);     % empty signal
stV = st0;
stV(fs+1 : fs+duration1*fs) = vis;  % visual flash starts at 1 s

%% Run MCD model on actual experimental trials
for iLag = 1:5
    for iDis = 1:4
        indA = round(data(iDis).responses(iLag,1) * fs);  % onset lag 
        stA = st0;
        stA(fs + indA : fs + indA + duration1*fs - 1) = data(iDis).signal';

        signals = [stV'; stA'];  % stack as [visual; auditory]
        [MCD_cor(iLag, iDis), MCD_lag(iLag, iDis)] = MCDq(signals, fs, param); % run MCD model
    end
end

%% Z-score normalize the MCD outputs
MCD_corMN = mean(MCD_cor(:));
MCD_corSD = std(MCD_cor(:));
MCD_lagMN = mean(MCD_lag(:));
MCD_lagSD = std(MCD_lag(:));

MCD_cor = (MCD_cor - MCD_corMN) / MCD_corSD;
MCD_lag = (MCD_lag - MCD_lagMN) / MCD_lagSD;

%% Fit probit regression to experimental responses using MCD predictions (Equation 10)
pred = [MCD_cor(:), MCD_lag(:)];
dat = vertcat(data.responses); dat = dat(:,2);  % binary responses

[b, ~, stats] = glmfit(pred, dat, 'binomial', 'link', 'probit');
yhat = glmval(b, pred, 'probit');

%% Run smooth sweep of lag values to generate psychometric functions
inds = 0:1:round(0.2 * fs);  % lag steps from 0 to 200 ms
nLag = numel(inds);
lags_plot = inds / fs;

for iLag = 1:nLag
    for iDis = 1:4
        indA = inds(iLag);  % lag in samples
        stA = st0;
        stA(fs + indA : fs + indA + duration1*fs - 1) = data(iDis).signal';

        signals = [stV'; stA'];
        [MCD_cor_plot(iLag, iDis), MCD_lag_plot(iLag, iDis)] = MCDq(signals, fs, param);
    end
end

% Normalize the predicted values using earlier computed means/SDs
MCD_cor_plot = (MCD_cor_plot - MCD_corMN) / MCD_corSD;
MCD_lag_plot = (MCD_lag_plot - MCD_lagMN) / MCD_lagSD;

%% Plotting
figure('Color', 'w');

for iDis = 1:4
    %% Get psychometric function (Equation 10)
    X = [MCD_cor_plot(:,iDis), MCD_lag_plot(:,iDis)];
    MCD_fit(:, iDis) = glmval(b, X, 'probit');

    % Find PSE (lag at which p(resp) ~ 0.5)
    [~, ind_pse] = min(abs(MCD_fit(:, iDis) - 0.5));
    pse(iDis) = lags_plot(ind_pse);

    %% Panel 1: Acoustic envelopes
    subplot(1,3,1); hold on; box on;
    plot((1:length(data(iDis).signal)) / fs, data(iDis).signal, ...
        'LineWidth', 1.2, 'DisplayName', [num2str(iDis*10) ' m']);
    xlabel('Time (s)');
    ylabel('Intensity');
    title('Acoustic Stimuli (envelopes)');
    set(gca, 'FontSize', 10);

    %% Panel 2: Psychometric curve
    subplot(1,3,2); hold on; box on;
    plot(lags_plot, MCD_fit(:, iDis), 'LineWidth', 2);
    scatter(data(iDis).responses(:,1), data(iDis).responses(:,2), ...
        40, 'filled');
    xlabel('Lag (s)');
    ylabel('p(Auditory First)');
    title('Psychometric Function');
    ylim([0 1]);
    set(gca, 'FontSize', 10);
end

legend(subplot(1,3,1), {'5m', '10m', '20m', '40m'}, 'Location', 'northeast');

%% Panel 3: PSS as a function of distance
subplot(1,3,3); hold on; box on;
plot([5, 10, 20, 40], pse, 'ko-', ...
    'LineWidth', 2, 'MarkerFaceColor', 'k');
xlabel('Distance (m)');
ylabel('PSS (s)');
title('PSS vs. Distance');
xlim([0 50]); ylim([0 0.2]);
set(gca, 'FontSize', 10);

sgtitle('MCD Model of Alais & Carlile (2005)', 'FontSize', 14, 'FontWeight', 'bold');
```
