## supplementary codes for "A Stimulus-Computable Model for Audiovisual Perception and Spatial Orienting in Mammals": MCDq.m

```
function [MCD_corr, MCD_lag, MCD_output] = MCDq(signals,fs,param)
%MCD computes the outputs of the MCD model
%
%   [MCD_corr, MCD_lag] = MCD(signals,fs) calculates the mean of the
%   output of an MCD unit (Equations 6 and 7, respectively) for audiovisual
%   time-varying signals. Signals is a 2-by-m matrix, whose first row
%   contains the visual signal, and the second row contains the auditory
%   signal. Fs is the sampling frequency of the signals (Hz).
%
%   [MCD_corr, MCD_lag] = MCD(signals,fs,param) sets the temporal constants
%   for the four tempoarl filters .
%
%   [MCD_corr, MCD_lag, MCD_output] = MCD(signals,fs) and
%   [MCD_corr, MCD_lag, MCD_output] = MCD(signals,fs,param)
%   additionally returns a structure with all filters, stimuli, and
%   time-varying model responses at the various steps of processing

%   Written by Cesare V. Parise (2023). 


nsamp=length(signals);
duration=nsamp/fs;
t=linspace(0,duration,nsamp);
dt=1/fs;
slow_n = 9;
fast_n = 6;

stimV=signals(1,:);
stimA=signals(2,:);

% Temporal constants
if nargin<3
    param=[0.0448821, 0.0369702, 0.180101, 0.180101];
end

% Generate filters
beta = 1;
% fv=((t./param(1)).* exp(-t./param(1)).*(1-beta*((t./param(1)).^2)/factorial(3)));
% fa=((t./param(2)).* exp(-t./param(2)).*(1-beta*((t./param(2)).^2)/factorial(3)));
fv1=((t./param(1)).^slow_n.*exp(-t./param(1)).*(1/factorial(slow_n)-beta*((t./param(1)).^2)/factorial(slow_n+2)));
fv2=((t./param(1)).^fast_n.*exp(-t./param(1)).*(1/factorial(fast_n)-beta*((t./param(1)).^2)/factorial(fast_n+2)));
fa1=((t./param(2)).^slow_n.*exp(-t./param(2)).*(1/factorial(slow_n)-beta*((t./param(2)).^2)/factorial(slow_n+2)));
fa2=((t./param(2)).^fast_n.*exp(-t./param(2)).*(1/factorial(fast_n)-beta*((t./param(2)).^2)/factorial(fast_n+2)));
 
beta = 0;
fva=((t./param(3)).* exp(-t./param(3)).*(1-beta*((t./param(3)).^2)/factorial(3)));
fav=((t./param(4)).* exp(-t./param(4)).*(1-beta*((t./param(4)).^2)/factorial(3)));

% early filtering
st_v1=dt*conv(fv1,stimV,'full');
st_v2=dt*conv(fv2,stimV,'full');
st_a1=dt*conv(fa1,stimA,'full');
st_a2=dt*conv(fa2,stimA,'full');

% squaring
st_v1=st_v1(1:length(t)).^2;
st_v2=st_v2(1:length(t)).^2;
st_a1=st_a1(1:length(t)).^2;
st_a2=st_a2(1:length(t)).^2;

% sum and sqrt
st_v = sqrt(st_v1+st_v2);
st_a = sqrt(st_a1+st_a2);

% late filtering
st_v_av=dt*conv(fva,st_v,'full');
st_a_va=dt*conv(fav,st_a,'full');

st_v_av=st_v_av(1:length(t));
st_a_va=st_a_va(1:length(t));

% xcorrelate
u1=st_a_va.*st_v;       
u2=st_v_av.*st_a;       

% MCD correlation detector output
% MCD_corr_signal=sqrt(u2.*u1);  
MCD_corr_signal=(u2.*u1);  
MCD_corr=mean(MCD_corr_signal); 

% MCD lag detector output
MCD_lag_signal=-u2+u1;         
MCD_lag=mean(MCD_lag_signal); 

if nargout==3
    MCD_output.corr = MCD_corr_signal;
    MCD_output.lag = MCD_lag_signal;
    MCD_output.v = signals(1,:);
    MCD_output.a = signals(2,:);
    MCD_output.st_v = st_v;
    MCD_output.st_a = st_a;
    MCD_output.st_v2 = st_v_av;
    MCD_output.st_a2 = st_a_va;
    MCD_output.u1 = u1;
    MCD_output.u2 = u2;
    MCD_output.filtVslow = fv1;
    MCD_output.filtVfast = fv2;
    MCD_output.filtAslow = fa1;
    MCD_output.filtAfast = fa2;
    MCD_output.filtV2 = fva;
    MCD_output.filtA2 = fav;
end
```
