## supplementary codes for "A Stimulus-Computable Model for Audiovisual Perception and Spatial Orienting in Mammals": MCD_simulation_Koerding_3_param.m

```
%% Simulation of Koerding et al. (2007) PLoS ONE
% This script simulates the AV localization task of Koerding (Figure 2C)
% Reproduces Figure 4F from Parise (2025) eLife.
% Körding, K. P., Beierholm, U., Ma, W. J., Quartz, S., Tenenbaum, J. B., & Shams, L. (2007). Causal inference in multisensory perception. PLoS one, 2(9), e943.
% 
% Parameters: bias, visual sigma, auditory sigma
%
% Author: Cesare Parise (2025)

clc; clear; close all;

%% Load Koerding et al. (2007) data
load data_CI % Columns: [Vloc, Aloc, Vresp, Aresp]

nPos = 5;
x = linspace(-10, 10, nPos);  % Discretized spatial responses

% Preallocate response counters
respV = zeros(6,6,6);  % Visual response histograms
respA = zeros(6,6,6);  % Auditory response histograms

% Accumulate response frequencies
for tt = 1:size(data_CI,1)
    if all(data_CI(tt,3:4) >= 0)
        indsV = 1 + [data_CI(tt,1), data_CI(tt,2), data_CI(tt,3)];
        indsA = 1 + [data_CI(tt,1), data_CI(tt,2), data_CI(tt,4)];

        respV(indsV(1), indsV(2), indsV(3)) = respV(indsV(1), indsV(2), indsV(3)) + 1;
        respA(indsA(1), indsA(2), indsA(3)) = respA(indsA(1), indsA(2), indsA(3)) + 1;
    end
end

% Normalize to get probabilities (excluding zero-response bins)
mnV = sum(respV(:,:,1:end),3);
mnA = sum(respA(:,:,1:end),3);
pV = respV ./ mnV;
pA = respA ./ mnA;
pV = pV(:,:,2:end);
pA = pA(:,:,2:end);

%% MCD model setup
nxTot = 40; xTot = linspace(-10, 10, nxTot);
params = [-3.4103, nan, 1.3471, 5.7034]; % bias, visual sigma, auditory sigma

nSamp = 500;
As = zeros(1,nSamp); As(100) = 1;
Vs = zeros(1,nSamp); Vs(100) = 1;

% Simulate MCD responses for all visual-auditory combinations
for gg = 2:6
    for rr = 2:6
        V1 = normpdf(xTot, x(gg-1), params(3));
        A1 = normpdf(xTot, x(rr-1), params(4));

        for tt = 1:nxTot
            a = As .* A1(tt);
            v = Vs .* V1(tt);
            signals = [v; a];
            [~, ~, MCD_output] = MCDq(signals, 500);

            mcdAxt(:,tt)  = MCD_output.st_a';
            mcdVxt(:,tt)  = MCD_output.st_v';
            mcdAVxt(:,tt) = MCD_output.corr';
        end

        % Normalize response distributions
        mcdAVx = mean(mcdAVxt); mcdAVx = mcdAVx / sum(mcdAVx);
        mcdAx  = mean(mcdAxt);  mcdAx  = mcdAx  / sum(mcdAx);
        mcdVx  = mean(mcdVxt);  mcdVx  = mcdVx  / sum(mcdVx);

        respAVmcd(gg,rr,:) = reshape(mcdAVx, [1,1,nxTot]);
        respAmcd(gg,rr,:)  = reshape(mcdAx,  [1,1,nxTot]);
        respVmcd(gg,rr,:)  = reshape(mcdVx,  [1,1,nxTot]);
        respmcd(gg,rr)     = sum(mcdAVxt(:));
    end
end

%% Discretize model responses to match behavioral response categories
cutoffs = [-inf, -7.5, -2.5, 2.5, 7.5, inf];
for gg = 2:6
    for rr = 2:6
        for tt = 1:5
            inds = xTot > cutoffs(tt) & xTot <= cutoffs(tt+1);
            respAVmcd2(gg,rr,tt) = sum(respAVmcd(gg,rr,inds));
            respAmcd2(gg,rr,tt)  = sum(respAmcd(gg,rr,inds));
            respVmcd2(gg,rr,tt)  = sum(respVmcd(gg,rr,inds));
        end

        % Normalize discretized responses
        respAVmcd2(gg,rr,:) = respAVmcd2(gg,rr,:) / sum(respAVmcd2(gg,rr,:));
        respAmcd2(gg,rr,:)  = respAmcd2(gg,rr,:)  / sum(respAmcd2(gg,rr,:));
        respVmcd2(gg,rr,:)  = respVmcd2(gg,rr,:)  / sum(respVmcd2(gg,rr,:));
    end
end

%% Compute mixture weight (Equation 19)
temp = respmcd(2:end, 2:end);
temp = (temp - mean(temp(:))) / std(temp(:));
respmcd(2:end, 2:end) = temp;

wAV = normcdf(respmcd + params(1));

% Combine responses 
respA2 = nan(size(pA));
respV2 = nan(size(pV));
for gg = 2:6
    for rr = 2:6
        for tt = 1:5
            respA2(gg,rr,tt) = wAV(gg,rr) * respAVmcd2(gg,rr,tt) + (1 - wAV(gg,rr)) * respAmcd2(gg,rr,tt);
            respV2(gg,rr,tt) = wAV(gg,rr) * respAVmcd2(gg,rr,tt) + (1 - wAV(gg,rr)) * respVmcd2(gg,rr,tt);
        end
    end
end

%% Display: MCD predictions (lines) vs data (dots)
mSz = 10;
figure(1); clf;
n = 0;
for gg = 2:6
    for rr = 2:6
        n = n + 1;
        v = squeeze(pV(gg,rr,:));
        a = squeeze(pA(gg,rr,:));
        avv = squeeze(respV2(gg,rr,:));
        ava = squeeze(respA2(gg,rr,:));

        subplot(5,5,n); hold on; box on;
        plot(x, v, '.b', 'MarkerSize', mSz);
        plot(x, a, '.r', 'MarkerSize', mSz);
        plot(x, avv, 'b', 'LineWidth', 1.5);
        plot(x, ava, 'r', 'LineWidth', 1.5);
        xlim([-10 10]); ylim([0 1]);
        xlabel('Location (deg)'); ylabel('p(resp)');
    end
end

%% Correlation between MCD and behavioral data
a = respA2(2:end,2:end,:);
b = pA(2:end,2:end,:);
c = respV2(2:end,2:end,:);
d = pV(2:end,2:end,:);

all_model = [a(:); c(:)];
all_data  = [b(:); d(:)];

corrMCDdata = corr(all_model, all_data);
```
