## supplementary codes for "A Stimulus-Computable Model for Audiovisual Perception and Spatial Orienting in Mammals": Run_MCD.m

```
% Run_MCD runs the MCD population model on a real-life movie
% The input video is the one used for Supplementary Video 3 and Figure 5
% Written by Cesare V. Parise (2025). 

clc; clear; close all

% load audiovisual stimuli
[aud, fsA] = audioread('Ventriloquist_Movie.mp4');
aud = mean(aud,2);

v = VideoReader('Ventriloquist_Movie.mp4');
fsV = v.FrameRate;

% get video frames, blur and rescale values to the interval [0,1]
nFr = 0; vid = []; vid2 = [];
while hasFrame(v)
    nFr = nFr+1;
    frame = readFrame(v);
    frame = double(rgb2gray(frame));

    vid(:,:,nFr) = frame;

    if  nFr==1
        frame1 = frame;
    else
        frame = frame-frame1;
    end

    vid2(:,:,nFr) = imgaussfilt(frame,3); % vary this to change the level of blur.

end
vid = rescale(vid);
vid2 = rescale(vid2);

% get audio envelope, resample and rescale values to the interval [0,1]
audEnv = envelope(aud,441,'rms');
tResampAud = linspace(0,size(vid,3)./fsV,size(vid,3));
tAud = linspace(0,numel(aud)./fsA,numel(aud));
ts = timeseries(audEnv,tAud);
ts = resample(ts,tResampAud);
audEnv = rescale(squeeze(ts.data));

% run MCD on video (pixel by pixel)
MCDcorr = nan(size(vid2));
MCDlag = nan(size(vid2));
for nRow = 1:size(vid,1)
    for nCol = 1:size(vid,2)
        signalV = vid2(nRow,nCol,:);
        signals = [signalV(:)';audEnv'];

        [~,~, MCD_output] = MCDq(signals,fsV);

        MCDcorr(nRow,nCol,:) = reshape(MCD_output.corr,[1,1,size(vid,3)]);
        MCDlag(nRow,nCol,:) = reshape(MCD_output.lag,[1,1,size(vid,3)]);
    end
end

% display results
MCDcorr = 1-rescale(MCDcorr);
MCDlag = rescale(MCDlag);

figure
for nFr = 1:size(vid,3)
    subplot(2,2,1)
    imshow(vid(:,:,nFr));
    title('Input video')
    
    subplot(2,2,3)
    cla; hold on
    plot(tResampAud, audEnv,'k')
    plot([tResampAud(nFr),tResampAud(nFr)],[0,1.1],'r')
    pbaspect([size(vid,2),size(vid,1),1]); set(gca,'YTick',[])
    xlim([0,tResampAud(end-1)]); ylim([0,1.1])
    title('Input audio'); xlabel('Time (s)'); ylabel('Amplitude')
    
    subplot(2,2,2)
    imshow(MCDcorr(:,:,nFr));
    title('Output MCD_{Corr}')
    
    subplot(2,2,4)
    imshow(MCDlag(:,:,nFr));
    title('Output MCD_{Lag}')
    
    drawnow
end
```
