## supplementary codes for "A Stimulus-Computable Model for Audiovisual Perception and Spatial Orienting in Mammals": MCD_Simulation_Mohl_et_al_2020.m

```
%% MCD modelling of Mohl 
% MCD simulation of gaze direction towards visual and acoustic cues in
% humans and monkeys
% Mohl, J. T., Pearson, J. M., & Groh, J. M. (2020). Monkeys and humans implement causal inference to simultaneously localize auditory and visual stimuli. Journal of Neurophysiology,

clc; clear; close all;

% Choose subject type: 1 = monkey, 0 = human
dataMonkey = 0;

% Load appropriate data and model parameters
if dataMonkey == 1
    load dataAM    % auditory unimodal (monkey)
    load dataVM    % visual unimodal (monkey)
    load dataAVM   % audio-visual bimodal (monkey)
    param = [-1.613, 3.4046, 0.1603]; % Equation 21: Beta_Crit, Beta_Corr, p_Lapse
else
    load dataAH    % auditory unimodal (human)
    load dataVH    % visual unimodal (human)
    load dataAVH   % audio-visual bimodal (human)
    param = [-5.5785, 7.4311, 0.0648]; % Equation 21: Beta_Crit, Beta_Corr, p_Lapse
end

%% Preprocess unimodal auditory responses
tA = unique(dataA(:,1));
for tt = 1:numel(tA)
    temp = dataA(dataA(:,1) == tA(tt), 2);
    matA(tt,:) = [tA(tt), mean(temp), std(temp)]; % unimodal mean and std
end

%% Preprocess unimodal visual responses
tV = unique(dataV(:,1));
for tt = 1:numel(tV)
    temp = dataV(dataV(:,1) == tV(tt), 2);
    matV(tt,:) = [tV(tt), mean(temp), std(temp)]; % unimodal mean and std
end

%% Compute empirical P(single saccade) as a function of AV disparity
dataAVc = dataAV(:,1) - dataAV(:,2);                % disparity
dataAVc(:,2) = dataAV(:,3) == dataAV(:,4);          % 1 if same saccade
dx = unique(dataAVc(:,1));
for tt = 1:numel(dx)
    CI(tt,:) = [dx(tt), mean(dataAVc(dataAVc(:,1) == dx(tt), 2))];
end

%% Histogram setup
nPos = 862;
edges = linspace(-31, 31, nPos + 1);
pos_hist = linspace(-30.5, 30.5, nPos);

% Prepare for looping over all audio-visual trial types
tA = unique(dataAV(:,2));
tV = unique(dataAV(:,1));
dataCell = {};
dataCell_plot = {};
n_plot = 0;
n = 0;

for iA = 1:numel(tA)
    for iV = 1:numel(tV)
        n = n + 1;

        % Extract auditory and visual responses
        rA = dataAV(dataAV(:,2)==tA(iA) & dataAV(:,1)==tV(iV), 4);
        rV = dataAV(dataAV(:,2)==tA(iA) & dataAV(:,1)==tV(iV), 3);

        % Same vs. different saccades
        indSame = (rA == rV);
        same = rA(indSame);
        difA = rA(~indSame);
        difV = rV(~indSame);

        % Build histograms without plotting
        dataCell{n}(1,:) = histogram(same, edges, 'Visible', 'off').Values;
        dataCell{n}(2,:) = histogram(difV, edges, 'Visible', 'off').Values;
        dataCell{n}(3,:) = histogram(difA, edges, 'Visible', 'off').Values;
        dataCell{n}(4,:) = histogram(rV, edges, 'Visible', 'off').Values;
        dataCell{n}(5,:) = histogram(rA, edges, 'Visible', 'off').Values;

        % Only proceed if there's data
        if sum(dataCell{n}(:)) > 0
            n_plot = n_plot + 1;
            dataCell_plot{n_plot} = dataCell{n};

            % Parameters for MCD model
            para = [tV(iV), tA(iA), mean(matV(:,3)), mean(matA(:,3))];
            MCD_CI(n_plot) = runMCD_CI(para);

            % Record positions and single-saccade proportion
            pos_plot(n_plot,:) = [tV(iV), tA(iA)];
            df(n_plot) = tA(iA) - tV(iV);
            ci(n_plot) = mean(indSame);
        end
    end
end

%% Normalize MCD outputs
for tt = 1:n_plot
    MCD_CI(tt).mcdVxtn = MCD_CI(tt).mcdVxt / sum(MCD_CI(tt).mcdVxt(:));
    MCD_CI(tt).mcdAxtn = MCD_CI(tt).mcdAxt / sum(MCD_CI(tt).mcdAxt(:));
    MCD_CI(tt).mcdAVxtn = MCD_CI(tt).mcdAVxt / sum(MCD_CI(tt).mcdAVxt(:));
end

%% Get P(single saccade)
for tt = 1:n_plot
    mcdci(tt) = MCD_CI(tt).respmcd;
end

for tt = 1:numel(dx)
    mcdCI0(tt) = log10(mean(mcdci(df == dx(tt))));
end

% Z-score normalization
mn1 = mean(mcdCI0);
sd1 = std(mcdCI0);
predictor = (log10(mcdci) - mn1) / sd1;

% p(single fixation) Equation 21
mcdci2 = normcdf(param(2) * predictor + param(1));
mcdci2 = param(3) + (mcdci2 * (1 - 2 * param(3)));

% Compute model predictions by disparity
for tt = 1:numel(dx)
    mcdCI(tt,:) = [dx(tt), mean(mcdci2(df == dx(tt)))];
end


% Compute disparities and sort
disparities = pos_plot(:,2) - pos_plot(:,1);  % A - V
[sorted_deltas, sort_idx] = sort(disparities);

figure(1); clf;
set(gcf, 'Color', 'w');
tiledlayout(2, 10, 'Padding', 'compact', 'TileSpacing', 'compact');

for plotIdx = 1:n_plot
    tt = sort_idx(plotIdx);

    % Compute tile index with zigzag logic
    if plotIdx <= 10
        row = 1;
        col = plotIdx;
    else
        row = 2;
        col = 20 - plotIdx + 1;  % Flip order on bottom row
    end
    tileIdx = (row - 1) * 10 + col;

    nexttile(tileIdx);
    hold on; box on;

    % MCD model mixture
    mcdV = sum(MCD_CI(tt).mcdVxtn);
    mcdA = sum(MCD_CI(tt).mcdAxtn);
    mcdAV = sum(MCD_CI(tt).mcdAVxtn);
    respV(tt,:) = (1 - mcdci2(tt)) * mcdV + mcdci2(tt) * mcdAV;
    respA(tt,:) = (1 - mcdci2(tt)) * mcdA + mcdci2(tt) * mcdAV;

    % Empirical data
    dtV = dataCell_plot{tt}(4,:) / sum(dataCell_plot{tt}(4,:));
    dtA = dataCell_plot{tt}(5,:) / sum(dataCell_plot{tt}(5,:));

    % Plot stimulus markers
    xline(pos_plot(tt,1), '-', 'Color', [0.4 0 0.8], 'LineWidth', 1.2);  % Visual
    xline(pos_plot(tt,2), '--', 'Color', [0 0.5 0], 'LineWidth', 1.2);   % Auditory

    % Plot response curves
    plot(pos_hist, smooth(dtV, 17) + smooth(dtA, 17), 'k', 'LineWidth', 1.2);  % Empirical
    plot(pos_hist, respV(tt,:) + respA(tt,:), 'b', 'LineWidth', 1.2);         % Model

    % Axes
    xlim([-30, 30]);
    ylim([0, 0.06]);
    set(gca, 'XTick', -30:15:30, 'YTick', []);
    title(['\Delta = ' num2str(disparities(tt)) '°'], 'FontSize', 9);
end

sgtitle('MCD vs Empirical Responses ', ...
    'FontSize', 14, 'FontWeight', 'bold');


figure(2); clf;
set(gcf, 'Color', 'w', 'Position', [500 300 500 400]);

% Plot empirical and model-predicted p(single fix)
hold on; box on; grid on;

plot(CI(:,1), CI(:,2), 'ko', 'MarkerSize', 6, 'LineWidth', 1.2, 'DisplayName', 'Empirical');
plot(mcdCI(:,1), mcdCI(:,2), 'b-', 'LineWidth', 2, 'DisplayName', 'Model');

ylim([0, 1]); xlim([-30, 30]);
xlabel('Disparity (deg)', 'FontSize', 12);
ylabel('P(Single Fixation)', 'FontSize', 12);
title('Causal Inference Fit', 'FontSize', 14, 'FontWeight', 'bold');
legend('Location', 'best');
```
