## supplementary codes for "A Stimulus-Computable Model for Audiovisual Perception and Spatial Orienting in Mammals": runMCD_CI.m

```
function MCD_CI = runMCD_CI(para)
% RUNMCD_CI Simulates causal inference using the MCD model
%
% INPUT:
%   para = [posV, posA, stdV, stdA]
%     posV : position of the visual stimulus
%     posA : position of the auditory stimulus
%     stdV : standard deviation of visual input
%     stdA : standard deviation of auditory input
%
% OUTPUT:
%   MCD_CI : structure containing simulated MCD outputs
%     .mcdVxt   : MCD trace for visual input
%     .mcdAxt   : MCD trace for auditory input
%     .mcdAVxt  : MCD correlation trace (multisensory)
%     .respmcd  : scalar summary statistic (total AV correlation)

% Unpack input
posV = para(1);
posA = para(2);
stdV = para(3);
stdA = para(4);

% Define position space
nPos = 862;
pos = linspace(-30.5, 30.5, nPos);  % Histogram bin centers

% Number of MCD samples (simulations)
nSamp = 1500;

% Preallocate output arrays
mcdAxt  = zeros(nSamp, nPos);  % MCD Auditory spacetime matrix
mcdVxt  = zeros(nSamp, nPos);  % MCD Visual spacetime matrix
mcdAVxt = zeros(nSamp, nPos);  % MCD AV spacetime matrix
Axt     = zeros(nSamp, nPos);  % Auditory stimuli spacetime matrix
Vxt     = zeros(nSamp, nPos);  % Visual stimuli spacetime matrix

% Create unimodal stimuli
V1 = normpdf(pos, posV, stdV);  % Visual stimuli (spatial profile)
A1 = normpdf(pos, posA, stdA);  % Auditory stimuli (spatial profile)

% Normalize
V1 = V1 / sum(V1);
A1 = A1 / sum(A1);

% Initial state (single unit active at index 1)
As = zeros(1, nSamp); As(100) = 1;
Vs = zeros(1, nSamp); Vs(100) = 1;

% Iterate over positions to simulate the temporal MCD response
for tt = 1:nPos
    % Stimulus drive at position tt for each sample
    a = As * A1(tt);  % Auditory signal strength
    v = Vs * V1(tt);  % Visual signal strength

    % Combine signals into input for MCD
    signals = [v; a];  % 2 x nSamp matrix

    % Call MCD model 
    [~, ~, MCD_output] = MCDq(signals, 250);

    % Extract and store time series data
    mcdAxt(:,tt)  = MCD_output.st_a';   % Auditory MCD response
    mcdVxt(:,tt)  = MCD_output.st_v';   % Visual MCD response
    mcdAVxt(:,tt) = MCD_output.corr';   % AV correlation response
    Axt(:,tt)     = MCD_output.a';      % Raw auditory signal
    Vxt(:,tt)     = MCD_output.v';      % Raw visual signal
end

% Compute a summary statistic: total integrated AV correlation
respmcd = sum(mcdAVxt(:));

% Package output into structure
MCD_CI.mcdVxt   = mcdVxt;
MCD_CI.mcdAxt   = mcdAxt;
MCD_CI.mcdAVxt  = mcdAVxt;
MCD_CI.respmcd  = respmcd;

end
```
